# Language Models for Molecular Dynamics

**DOI:** 10.1101/2024.11.25.625337

**Authors:** Mhd Hussein Murtada, Z. Faidon Brotzakis, Michele Vendruscolo

## Abstract

Molecular Dynamics (MD) simulations provide accurate descriptions of the motions of molecular systems, yet their computational demands pose significant challenges in applications in molecular biology and materials science. Given the success of deep learning methods in a wide range of fields, a timely question concerns whether these methods could be leveraged to improve the efficiency of MD simulations. To investigate this possibility, we introduce Molecular Dynamics Language Models (MDLMs), to enable the generation of MD trajectories. In the present implementation, an MDLM is trained on a short classical MD trajectory of a protein, where structural accuracy is maintained through kernel density estimations derived from extensive MD datasets. We illustrate the application of this MDLM in the case of the determination of the free energy landscape a small protein, showing that this approach makes it possible to discover conformational states undersampled in the training data. These results provide initial evidence for the use of language models for the efficient implementation of molecular dynamics.

## 1 Introduction

Proteins are dynamic molecules whose function is intimately linked with their ability to sample different conformational states [14, 6, 7, 26, 43]. Since protein motions underlie most biological processes, the ability to characterise them is crucial in a wide range of applications [14, 6, 7, 26, 43].

As many functional motions of proteins involve the exploration of the conformational space under equilibrium conditions, several approaches have focused on the generation of conformational ensembles obeying the Boltzmann distribution [10, 8]. Based on the remarkable success of machine learning (ML) methods in protein structure predictions [19, 1, 21], one can ask whether the exploration of the Boltzmann ensembles of proteins could be implemented using some forms of ML [37, 28, 17, 41, 18, 2].

In many applications, however, one is interested in following the protein dynamics according to the physical laws of motion. Over the last 60 years, molecular dynamics (MD) simulations have represented the gold standard for this purpose, as they provide atomic-level detail by implementing the exploration of the conformational space of proteins by integrating numerically the equations of motion [32, 25, 20]. However, achieving convergence in MD simulations often requires extensive computational resources and time [32], or the application of system-specific enhanced sampling methods, particularly when exploring rare events or high-energy states [38, 36, 23]. This computational burden significantly limits the current ability to study long-timescale processes and rare conformational transitions. The challenge lies in combining the efficiency of ML with the physical accuracy of MD simulations. A successful approach ought to balance multiple competing demands: adherence to physical laws, achievement of computational efficiency and accuracy in capturing conformational diversity [5, 4]. One way to address this problem involves methods based on the use of ML to speed up the calculations of force fields [4, 3, 29].

Language Models (LMs) offer opportunities to address the challenges involved in generating MD trajectories that capture the underlying laws of motion, as they are powerful tools for pattern recognition and generation [9]. Broadly speaking, by representing proteins as words and MD trajectories as sentences, one can ask whether LMs can capture the grammar - i.e. the physical laws - of protein motions.

Here, to investigate such opportunities, we describe Molecular Dynamics Language Models (MDLMs), a framework that bridges the physical accuracy of MD simulations with the pattern recognition capabilities of LMs. Our approach combines system-specific learning with broad physical principles. By training on a small fraction of typically required MD simulation time for a specific system, while incorporating physical insights derived from extensive MD datasets, MDLMs achieve both efficiency and accuracy. This physical guidance is implemented through kernel density estimations (KDEs) that capture allowed conformational regions based on 1390 MD trajectories from the ATLAS database [40]. We demonstrate the application of this approach in the case of the small protein chignolin [15] as a model system, where the MDLM discovers states not present or undersampled in the training data, while requiring only modest computational resources.

## 2 Methods

### 2.1 Overview of the methodological framework

The present implementation of the MDLM approach consists of three integrated components:

1. **System-specific learning from a short MD trajectory (up to 5% of a converged trajectory, referred to as the reference trajectory):** to train the LM, we run an MD simulation for a short period of time (equivalent to 5% of the reference trajectory) to generate the training data. For each frame, we calculate the backbone angle pairs (*ϕ*-*ψ*) and tokenize them to build a sentence describing the protein conformation at this frame. The final training dataset is a sequence of sentences.
2. **Physical guidance derived from extensive MD datasets (ATLAS database of MD simulations):** to ensure that the MDLM adheres to physical laws, we built an extensive dataset of neighbour-dependent Ramachandran distributions based on the MD simulations in the ATLAS database. We encode them as kernel density estimations (KDEs) and we use these KDEs as regularizes in the training loss, and as energy function in the sampling.
3. **Sampling process for reconstructing free energy landscapes:** after the LM model is trained, we run a sampling process where we use the LM model conditional inference to generate informed proposals. This enables efficient exploration of both low and high-energy states, while simultaneously reconstructing the free energy landscape of the reference MD simulation.

### 2.2 Structural Representation

While LMs operate in the continuous space during training and inference, they expect tokens (words) in their training data. Hence in the present MDLM, we tokenise the backbone angle pairs and represent protein conformations as sequences of tokens. This process makes MD data suitable for language models while preserving the essential physical characteristics of the protein structure. For a protein of length *N*, at each time step *t* in the training MD simulation, we represent the conformation as (*N* − 1) tokens (*τ*) plus a special fullstop token to separate time steps in the dataset: *S*_*t*_ = {*τ*_1_, *τ*_2_, …, *τ*_*N* − 1_,}. Each token *τ*_*i*_ encodes information about two neighbour residues and the angle pair in between: *τ*_*i*_ = (*r*_*i*_, *b*_*ϕ*_, *b*_*ψ*_, *r*_*i*+1_) where:

- *r*_*i*_, *r*_*i*+1_ ∈ 𝒜 (amino acid alphabet: A,C,D,E,F,G,H,I,K,L,M,N,P,Q,R,S,T,V,W,Y)
- 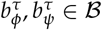 which is the set of letters a,b,c,… used to encode angle bins
- |ℬ| = *k* is the number of bins (e.g., if k=15, then ℬ = a,b,c,d,e,f,g,h,i,j,k,l,m,n,o)
- ‘.’ represents the special fullstop token separating time steps

For phi and psi, we use K-means clustering to calculate optimal **k** bin centres and then discretise the angles into these **k** bins:

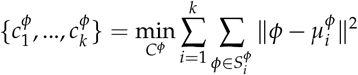

and

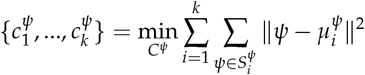

where:

- 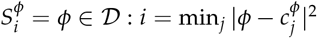
- 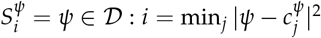
- 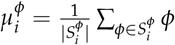
- 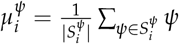

This adaptive K-means-based binning approach provides better alignment with the natural uneven distribution in Ramachandran plots [35]. So, regions that are more occupied by protein conformations are represented with finer granularity while less occupied regions are assigned to broader bins. This adaptive discretization preserves the natural distribution of protein conformations while enabling more accurate decoding of generated conformations (during inference).

- For encoding, at each time step *t*, we map the continuous angles to their corresponding letter bins by finding the nearest centroid. Let *θ* ∈ *ϕ, ψ*:

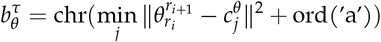

where *chr* and *ord* are character conversion functions that map between indices and ASCII letters. For example, if the nearest centroid is index 0, the angle would be encoded as ‘a’, if it’s index 1, it would be encoded as ‘b’, and so on.
- For decoding, we recover the continuous angles by mapping back from the letter bins to their corresponding centroid values:

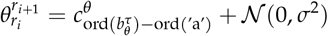

### 2.3 Physical Guidance through KDEs

The adherence to physical principles for the generated conformations in MDLMs is ensured through KDEs [11] derived from extensive MD data. ATLAS is an extensive database that gathers all-atom MD trajectories of 1390 protein systems, each run for 100 ns under consistent conditions (300 K, CHARMM36m force field) [40, 16]. We calculate *ϕ*-*ψ* pairs at each frame of these trajectories and generate a dataset of 400 neighbour-dependent Ramachandran distributions encoded as Kernel Density Estimations (KDEs) for every possible residue pair.

For any pair of amino acids (AA1, AA2), the corresponding KDE captures the joint probability distribution of their connecting *ϕ* and *ψ* angles:

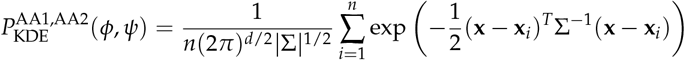

where:

- **x** = [*ϕ, ψ*]^*T*^ is the angle pair being evaluated
- **x**_*i*_ = [*ϕ*_*i*_, *ψ*_*i*_]^*T*^ are the observed angle pairs from MD
- Σ is the covariance matrix
- *d* = 2 is the dimensionality
- *n* is the number of observations
- angles are normalized by a scaling factor for numerical stability

These KDEs are used primarily to help the model adhere to the physical principles as they can be viewed as specialized Ramachandran plots that capture the conformational preferences specific to each amino acid pair, and hence during training and inference, they would guide the model to allowed, disallowed, favourable, and unfavourable regions. The probability densities encoded in these KDEs are:

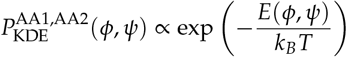

where *k*_*B*_ is the Boltzmann constant and *T* = 300*K* is the temperature at which the MD simulations were performed [40]. This allows us to interpret the negative log of the KDE as a free energy landscape:

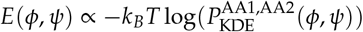

This relationship reveals that high-probability regions in the KDEs correspond to low free energy conformational states, while low-probability regions represent high free energy or forbidden conformations. This physical interpretation allows our framework to distinguish between favorable and unfavorable conformational changes, guiding the model towards physically realistic structures.

These free energy landscapes are used for inference and they capture key physical features:

- Primary minima corresponding to highly populated conformations
- Free energy barriers that determine conformational transition rates
- Amino acid specific preferences (e.g., the larger range of glycine and the narrower range of proline)
- Neighbor-dependent effects on backbone flexibility

### 2.4 Implementation Details and Training Setup

The present MDLM implementation is designed to run on consumer-grade system requirements, making it practical and accessible for widespread use in the research community.

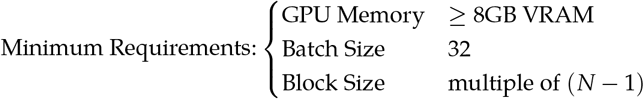

While these parameters ensure accessibility, the performance of the LM could potentially be enhanced by:

- Tuning the LM dimensions (*n*_embd_)
- Increasing batch sizes for better gradient estimation
- Extending block size for longer temporal dependencies in MD trajectories
- Adding additional attention heads to account for the complex patterns in MD

#### Training Data Selection

We select the initial 5% of a converged MD simulation of chignolin [24], which is composed of 10 amino acid residues (GYDPETGTWG). From the 5%, we mask out some high-energy and transition state structures. The structures were then tokenised as described in the Structural Representation section earlier.

We aim by this data selection approach to validate: (i) efficient learning from limited MD data, (ii) conformational diversity, (iii) coverage of high-energy states, and (iv) the ability of the model to recover rare conformations.

### 2.5 Language Model Architecture

The LM architecture adapted in the current implementation of MDLMs is the GPT-J architecture [44, 12] which we adapt for our goal of protein conformational generation. Mainly, we adjusted its default loss function to adhere to the physics of protein structures while simultaneously learning the temporal evolution of conformations in the training MD data.

#### Core Architecture

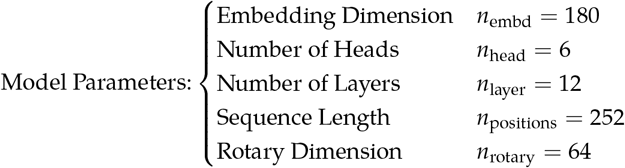

#### Rotary Positional Embeddings

The GPT-J model encodes positional information as rotary positional embeddings (RoPE) [39] which means that it introduces a rotational mechanism in its embedding space. This representation provides some advantages over traditional positional encoding, particularly for complex patterns such as conformational changes over time in MD:

- It enables the model to understand relative positions of angle pairs in protein sequences and in time-lagged trajectories
- It enhances the ability of the model to capture long range dependencies
- It is suitable for our periodic angle representation

#### Sequence Length

The sequence length of 252 allows processing multiple time steps together, where each time step is a sequence of length N-1. This enables the multi-head attention mechanism [42] to capture long range dependencies:

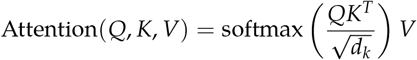

where Q, K, V are matrices incorporating both spatial (residue position) and temporal (time step) information. This mechanism allows the model to learn the patterns and correlations between different angle pairs of the protein representation across time. Additionally, it enables capturing both local and non-local conformational dependencies, while understanding temporal patterns in conformational changes.

#### Loss function

The loss function is designed to balance between accurate prediction of stable conformations and effective sampling of transition (high energy) states by adding a physics-based regularizer term based on the KDEs calculated from the MD database:

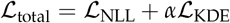

The first term, L_NLL_, is the standard next-token prediction negative log-likelihood:

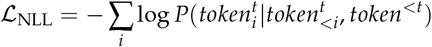

where the model learns to predict each token given both the spatial context (previous residues in the same time step) and temporal context (previous time steps).

The second term, ℒ_KDE_, is a weighted entropy term designed to prevent bias against high-energy conformations which naturally appear less in the training dataset:

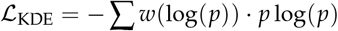

with Gaussian weight function:

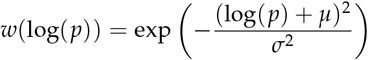

Properties of this weighting function:

1. It peaks at log(*p*) = −*µ*, which corresponds to medium-probability (allowed) regions in the conformational landscape
2. The width parameter (*σ*) is to determine the range of probabilities that receive significant weight
3. High-probability (low energy, stable) states are downweighted in order to prevent the model from overfitting to stable conformations
4. Very low-probability (physically forbidden) states receive minimal weight, maintaining physical feasibility

For chignolin, we set *µ* = 8 and *σ* = 4 which were determined through looking at the KDE probability distribution of angle pairs in the training data.

### 2.6 Inference

The conformation generation process in MDLMs comprises:

- LM inference: A beam search with multinomial sampling, conditional on random states from the 5% MD trajectory, predicts the next state in the form of MD trajectory starting from the random state conditioned on.
- Convergence acceleration: The predicted next state is penalized if the energy is increased significantly.

#### LM inference

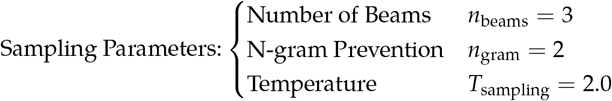

For each position *i*, the model output is a probability distribution over possible tokens in vocabulary:

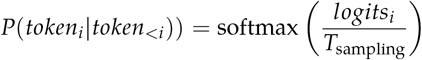

where *T*_sampling_ = 2.0 scales the logits to control sampling diversity. The beam search maintains the top-k partial sequences by score:

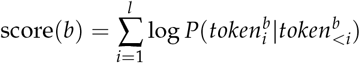

Advantages of parameters choice:

- Beam search with multinomial sampling allows for efficient exploration of paths with different probabilities and maintaining diverse candidates (stable and high energy conformations) [27] balancing exploration and exploitation.
- Elevated temperature (*T*_sampling_ *>* 1) flattens the distribution and encourages exploration of high energy regions [31]
- N-gram blocking prevents repetitive patterns which encourages diversity and creativity [45]

#### Convergence acceleration

The predicted next state is penalized to disfavor the generation of high-energy states:

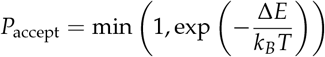

and

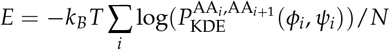

The physical temperature is set to T = 300K to match the MD training conditions, ensuring consistent energy scaling between the training data and sampling process.

## 3 Results

### 3.1 Ramachandran Analysis

We evaluate the ability of the current MDLM implementation to explore the conformational space of proteins through a Ramachandran analysis (Figures 5-6). For each peptide pair, we compare three distributions: the 5% training data, the MDLM-based sampling, and the reference (ground truth) from the complete MD trajectory. The overlap between sampled and reference conformational spaces is quantified using the Jaccard index (𝒥) [13], which ranges from 0 (no overlap) to 1 (perfect overlap). We calculate two (𝒥) scores for each residue pair along the amino acid sequence: one comparing the 5% training data with the full trajectory (𝒥_5_), and another comparing the MDLM sampling with the full trajectory (𝒥_*LM*_).

This MDLM achieves improvements in conformational space coverage for most residue pairs. For instance, the PE pair achieves significant enhancement in sampling, with 𝒥_5_=0.51 improving to 𝒥_*LM*_=0.82, indicating effective exploration of the conformational landscape from limited training data. Similarly, the GT pair shows an increase from 𝒥_5_=0.68 to 𝒥_*LM*_=0.94, effectively recovering conformational states that were poorly sampled or entirely missing in the training data.

Despite these improvements, the current MDLM exhibits two distinct types of sampling challenges:

- **Population accuracy:** The DP pair (𝒥_5_=0.58 → 𝒥_*LM*_=0.67) illustrates a case where the model discovers a conformational state around (−*π*/2, 0) that was completely absent in the 5% training data. While this finding demonstrates the MDLM ability to explore beyond its training distribution, it overestimates the population of this state compared to the full trajectory.
- **Additional populations:** The GY pair represents the only case where sampling quality decreases (𝒥_5_=0.88 → 𝒥_*LM*_=0.79), with the model exploring conformational regions not present in the full trajectory. This finding underscores the special conformational properties of glycine [30], and prompts further investigations on whether these additional populations were not sampled in the reference trajectory or were oversampled by the MDLM.

### 3.2 Reconstruction of the Free Energy Landscape

We next analyzed the ability of the current MDLM to reconstruct the complete free energy landscape using dimensionality reduction and KDE (Figure 2). The free energy landscapes are constructed by first converting the backbone angles (φ, ψ) to periodic features using sine and cosine transformations. These features undergo standardization followed by Independent Component Analysis (ICA) to obtain two time-independent components (tICs). The free energy at each point is calculated using:

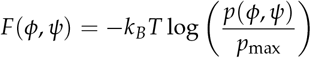

where *p*(*x*) is estimated using Gaussian kernel density estimation with bandwidth 0.08, and *T* = 300*K*. The resulting free energy landscapes are normalized relative to their global minimum.

**Figure 1:**
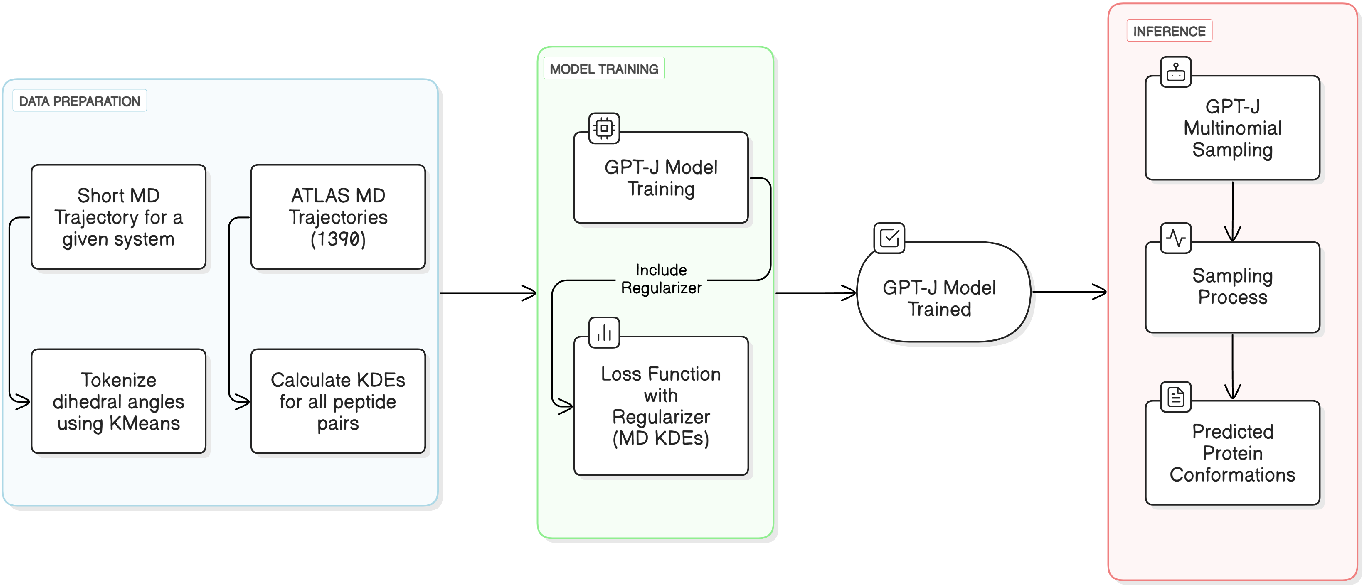
Flow diagram of the implementation of the MDLM approach discussed in this work.

**Figure 2:**
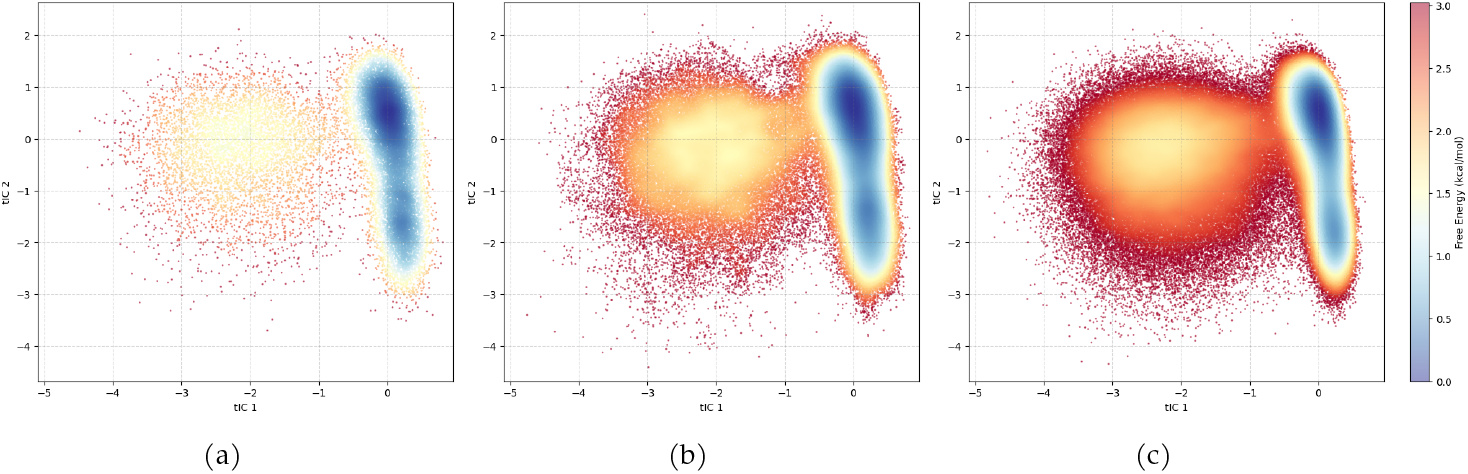
Comparison of the free energy landscapes for the training data (a), MDLM results (b), and reference trajectory (c).

Figure 2 reveals three distinct basins: two folded states on the right side of the landscape (*G*_1_, *G*_2_) and one unfolded state (U) on the left. Our sampling effectively recovers:

- Relative depths of the basins, with ΔG ≈ 2.5kcal/mol
- Inter-basin distances in tIC space matching literature values [24]
- The connectivity between states, including transition regions

At a structural level, the MDLM reproduces key structural features, including the characteristic turn geometry in the folded states and the extended conformations in the unfolded ensemble. Figure (3) shows representative structures from the sampling compared to those from equivalent basins in the reference trajectory, while figure (4) shows a sample of conformations generated by the framework which closely match unseen structures from the latter 95% of the trajectory.

**Figure 3:**
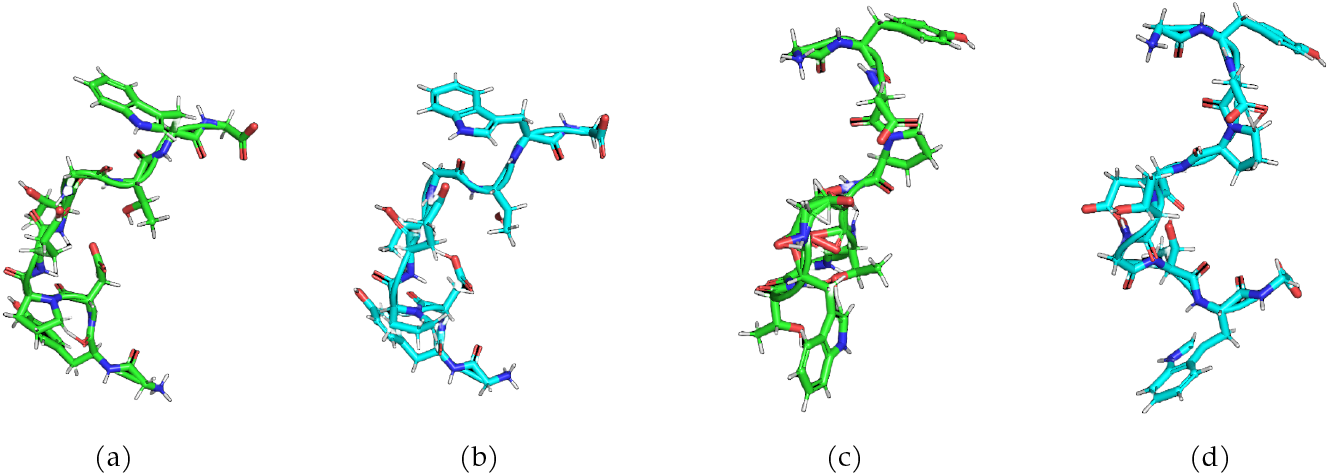
Comparison of folded and unfolded states from the reference and the MDLM simulations. **(a, b)**: Folded state at basin (0.18, 0.43) [RMSD= 1.69]. **(c, d)** Unfolded state at basin (−2.75, -0.34) [RMSD= 3.1]. MDLM conformations are shown in green and reference conformations in blue.

**Figure 4:**
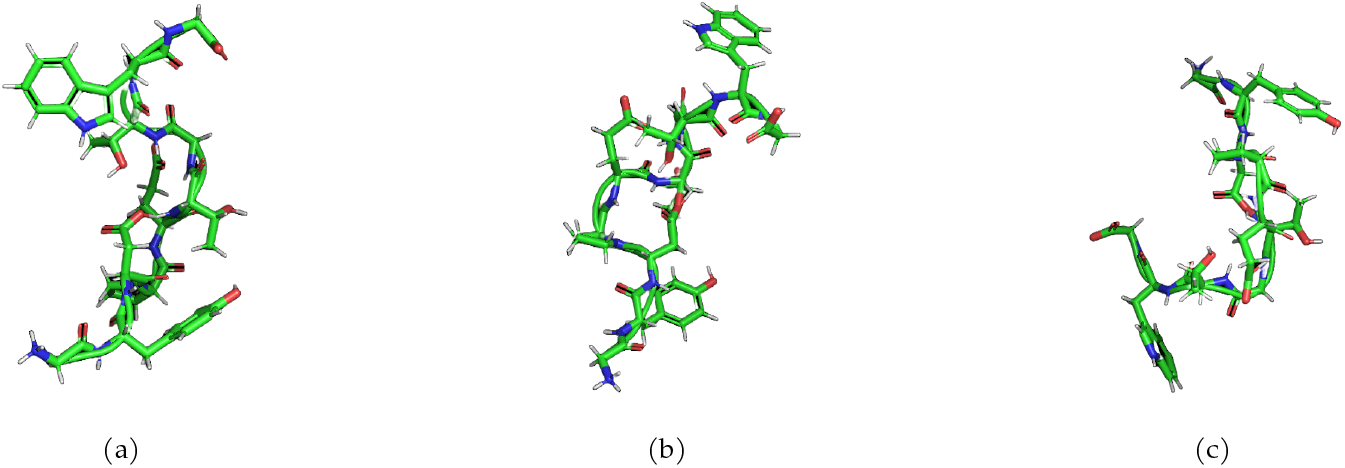
MDLM-generated conformations closely matching structures that only appear in the unseen 95% of the trajectory, were identified using k-nearest neighbors distance in the *ϕ*/*ψ* angle space. RMSDs ((a) 0.28, (b) 0.49, and (c) 0.38) were then calculated between matching structures using PyMol software.

**Figure 5:**
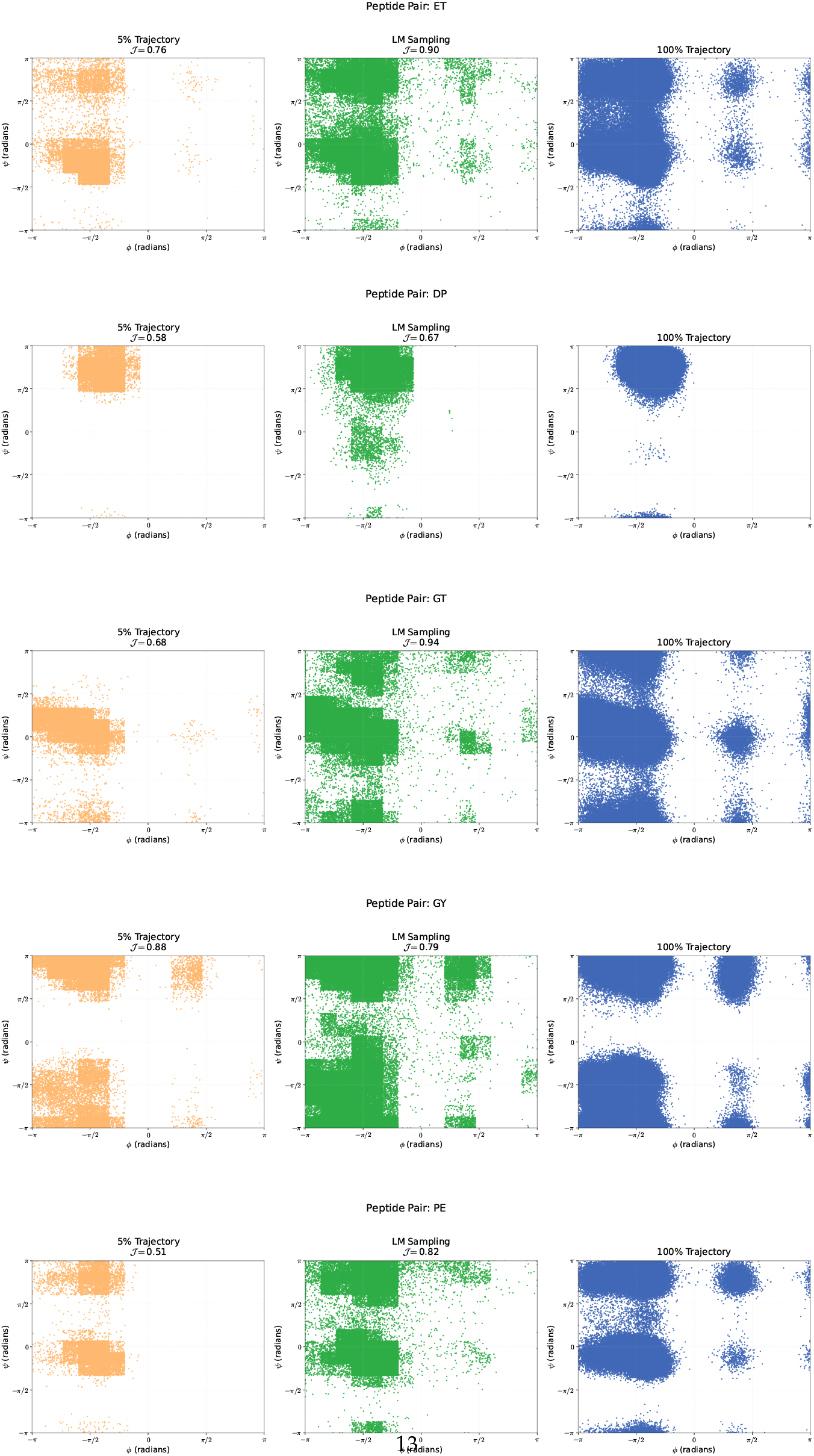
Ramachandran plots comparison for pairs: ET, DP, GT, GY, and PE

**Figure 6:**
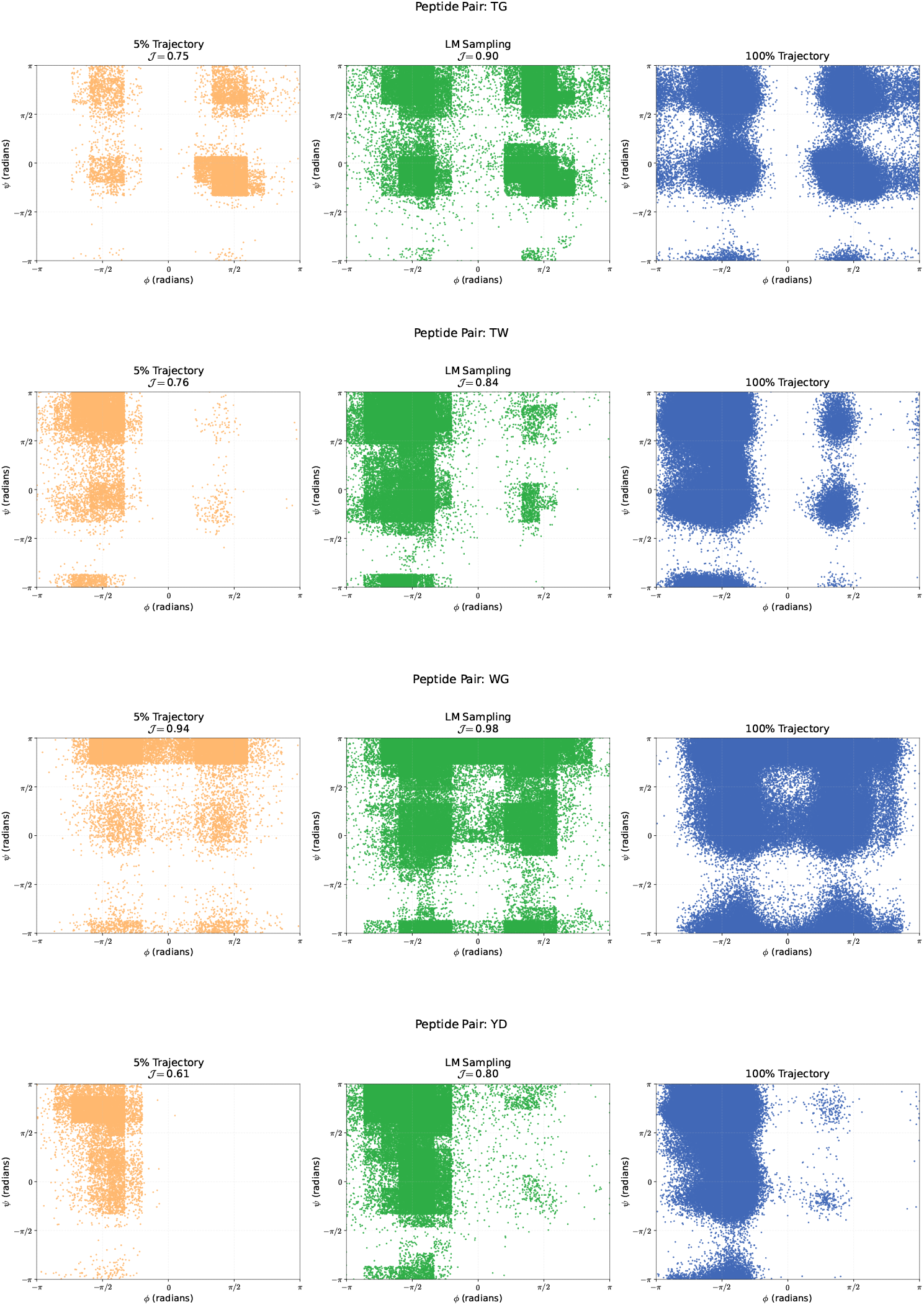
Ramachandran plots comparison for pairs: TG, TW, WG, YD

The recovery of the free landscape underlines the MDLM ability to capture both thermodynamic (basin depths) and kinetic (transition regions) features of the free energy landscape, despite training on only 5% of the trajectory data. Notably, the recovery is achieved using only 14 hours of combined CPU/GPU time on accessible hardware (RTX-4060 GPU, i7-12700H CPU), compared to the weeks of computation on specialized clusters typically required for full MD trajectories. The model training consumed 6 hours of GPU time, while the remaining 8 hours were split between GPU-accelerated sampling and CPU-based evaluation.

We note that the computation time to reach convergence is highly dependent on the size and quality of the training data, as it is used both for training the model and as the initial state for sampling.

### 3.3 Results Summary

The implementation of MDLMs that we reported demonstrates the ability to efficiently explore the conformational space of a protein while maintaining physical accuracy. Starting from just 5% of MD trajectory data, it achieves significant improvements in conformational sampling, with Jaccard indices improving in challenging cases like the PE residues pair. The reconstructed free energy landscape captures key thermodynamic features including the correct folded-unfolded energy difference (1.5 kcal/mol) and transition barriers (2.0 kcal/mol), aligning well with full trajectory. While some challenging cases exist, particularly for highly flexible sequences like GY, this MDLM demonstrates robust performance across different peptide pairs using only modest computational resources.

This work, focused on backbone conformations, serves as a proof of concept for MDLMs. Future development will extend to all-atom representation for enhanced accuracy and better handling of side-chain interactions. Additionally, optimization of the sampling process and physical guidance could further improve performance, particularly for challenging cases like highly flexible residues.

## 4 Discussion

The application of deep learning methods to structural biology has led to transformative approaches to protein structure determination, as it has become possible to predict the structures of proteins based on their amino acid sequences with an accuracy typically compared with that of high-resolution experimental methods [19, 1, 21]. Encouraging results have been obtained recently towards the use of deep learning methods for the description of the conformational space of proteins in terms of Boltzmann ensembles [37, 28, 17, 41, 18, 2]. A problem that remains open, however, concerns the use of deep learning methods for the prediction of protein dynamics.

Language models (LMs), particularly those based on the transformer architecture, such as the generative pre-trained transformer (GPT) series [34], have introduced innovative features that make them especially suitable for molecular conformational analysis. Evolutionary Scale Modeling (ESM) approaches have shown promise for protein structure prediction and design and for modeling protein evolution [22]. The causal nature of GPT models, where each token is generated based on all previous tokens, aligns naturally with the sequential nature of conformational changes in proteins. This causality ensures that each predicted structural element considers the entire preceding structural context, similar to how protein conformational changes propagate through the structure. The rotary positional embeddings in language models provide a powerful mechanism for understanding relative positions in sequences, allowing the model to effectively process and generate conformational changes while maintaining structural coherence [39, 33]. This feature is particularly valuable when exploring protein conformations, as it enables the model to swap structural elements while preserving their physical relationships. Furthermore, the beam search multinomial sampling strategy used in language models inference is crucial for generating diverse yet physically plausible conformations, as it allows for controlled exploration of the conformational space by maintaining multiple probable pathways during generation [27].

We note that other structural representations could be investigated, in addition the those used here in terms of dihedral angles. We are currently exploring the use of variational autoencoders [46] towards this goal.

In perspective, we would expect language models to be able to learn physical conservation rules from MD trajectories generated by laws of motion that obey such rules. In the present implementation, MDLMs are trained in a system-specific manner by using a small portion of a MD simulation of the protein of interest. Upon further developments, one can imagine training large language models (LLMs) on a large set of MD trajectories to establish procedures to run LLM-based MD simulations starting from the only knowledge of the amino acid sequence of the protein under study.

